# Complement receptor-mediated uptake into renal mononuclear phagocytes promotes intracellular UPEC persistence and limits β-lactam efficacy in pyelonephritis

**DOI:** 10.64898/2026.07.09.737431

**Authors:** Beata Goldyn, Olena Babyak, Pranitha Rokkam, Yevheniia Minchuk, Marigona Sutaj, Clivia Lisowski, Nicolette Bluszcz, Daniela Klaus, Annika Völkel, Amina Abdelmageed, Junping Yin, Andrew Kueh, Huiyun Hu, Isabelle Müller, Marco Herold, Sibylle von Vietinghoff, Daniel R. Engel, Natalio Garbi, Ulrich Dobrindt, Thomas Miethke, Florian Wagenlehner, Selina K. Jorch, Christian Kurts

## Abstract

**Introduction:** Acute pyelonephritis remains a major clinical problem. Relapses occur despite apparently appropriate antibiotic therapy, suggesting that uropathogenic Escherichia coli (UPEC) persist in intrarenal niches. Intracellular bacterial reservoirs are a plausible explanation, but the relevant host cells, entry mechanisms and therapeutic implications in the kidney remain undefined. In principle, such reservoirs should favor choosing intracellularly active antibiotics, but increasing resistance to many of these agents leaves β-lactams widely used in clinical practice, despite their predominantly extracellular activity.

**Methods:** We analyzed murine pyelonephritis to identify the cellular reservoir of persistent UPEC. We generated mice genetically deficient for complement receptors CR3 and CR4 and tested their role in bacterial entry and persistence *in vivo*. Pharmacological complement receptor inhibition was applied to assess whether blocking bacterial re-entry into host cells improves antibiotic efficacy.

**Results:** Renal MNP were identified as the major intracellular reservoir for UPEC in mice. Complement opsonization enabled bacterial entry into these cells through CR3 and CR4, allowing UPEC to evade neutrophil-mediated killing and extracellularly active antibiotics. Genetic deletion of CR3 and CR4 abolished intracellular bacterial persistence and reduced renal bacterial burden. Because MNP undergo physiological turnover, intracellular UPEC must periodically exit host cells and infect new ones. Pharmacological inhibition of complement receptors prevented such bacterial re-entry and enhanced the efficacy of β-lactam antibiotics which cannot penetrate cell membranes.

**Conclusions:** Complement receptor-mediated entry into renal MNP establishes an intracellular UPEC reservoir that promotes persistence during experimental pyelonephritis. Blocking these receptors prevents renewal of the intracellular niche and improves β-lactam efficacy in mice.

## INTRODUCTION

Urinary tract infections (UTIs) affect an estimated 150 million individuals worldwide each year^1^; 40-50% of women will experience a UTI over their lifetime^2^. UTIs affecting the bladder cause cystitis, while ascension to the kidneys results in pyelonephritis, a painful and serious complication that threatens renal function^3,4^ and increases the risk for bladder cancer^5^. While adjunct therapies exist for bacterial cystitis, treatment of pyelonephritis relies on antibiotics, which is an escalating clinical challenge due to increasing antibiotic resistance^6,7^.

Uropathogenic *Escherichia coli* (UPEC) cause over 80% of UTI^8^. The seminal discovery that these bacteria invade bladder epithelial cell^9^ refuted the longstanding belief that they are exclusively extracellular pathogens^10^. Some UPEC replicate in the cytosol of bladder cells, which is nutrient-rich and affords escape from immune surveillance. Alternatively, they may reside and replicate within host vacuoles^11^ or form biofilms^12,13^. This ability to “hide” within host epithelial cells is thought to explain relapses of UTIs and other bacterial infections, but a causal remains to be established^14^.

This ability may theoretically contribute also to the limited long-term effectiveness of antibiotic therapy^8,15^. β-lactams including penicillins and cephalosporins are frequently used for pyelonephritis but do not penetrate cell membranes. In current clinical practice, antibiotic choice is guided largely by disease severity and susceptibility testing in the setting of the world-wide rise of antimicrobial resistance^16–18^. Their ability to reach bacteria within host cells is rarely considered, although this property may critically influence treatment success if UPEC can hide intracellularly.

UPEC use uroplakin, a specialized transmembrane protein, to enter bladder uroepithelial cells^19,20^. UPEC establish intracellular niches also in the kidney during pyelonephritis, but renal parenchyma does not contain uroplakin^+^ epithelial cells^21–23^. Complement secreted by tubular epithelial cells (TECs) has been reported to be critical for establishing colonizing the kidney^24^, but these cells do not express receptors for complement opsonins either^21–23^, leaving the identity of the renal cellular reservoir unresolved.

Complement opsonization of microbes occurs by covalently attaching the C3b fragment to their surface for recognition by the complement receptors CR3 and CR4^25^. Both are heterodimers containing the β2 integrin, CD18, and either the α integrin CD11b (encoded by *Itgam*) in CR3, or CD11c (encoded by *Itgax*) in CR4. Mononuclear phagocytes (MNPs), including macrophages and dendritic cells, usually express CR3 and/or CR4. Some bacteria enter and hide within MNPs in a complement-dependent manner, such as *Mycobacterium* or *Salmonella*. MNPs exist in various tissue-specific subtypes and typically turn over by local proliferation or by replacement with bone marrow-derived precursors. Kidney MNPs are especially heterogenous and are replaced during adulthood every 2 weeks^26^. They serve as sentinel cells that detect pathogens or tissue damage and activate immune effector cells to combat microbes^27^. During pyelonephritis, MNPs sense UPEC and secrete chemokines that recruit neutrophils within hours^28,29^. Neutrophils are the principal immune effector cells against pyelonephritis^30,31^. They express CD11b, but typically lack CD11c.

Here, we aimed to identify the kidney cells invaded by UPEC during pyelonephritis and the molecular mechanisms involved. We demonstrate that renal MNP, and not epithelial cells, are the reservoir for relapsing pyelonephritis, and identify the entry receptors. This allowed us to conceptualize the first host-directed adjunct therapy against pyelonephritis.

## RESULTS

### UPEC hide within renal mononuclear phagocytes

Given that uroepithelial cells provide an intracellular niche for UPEC bacterial cystitis^9,19,20^, it is often assumed that epithelial cells provide a comparable niche also in pyelonephritis^24^. However, when we assessed the ability of UPEC to invade human HK-2 kidney tubular epithelial cells (TEC) and murine primary kidney epithelial cells *in vitro*, very low intracellular colonization was observed (Fig. 1A-D, Suppl. Video 1), much lower than did bladder uroepithelial cell (Fig. 1A, S1A). In these experiments, we had included bone-marrow-derived macrophages as a control, and noted effective intracellular entry of UPEC, but only when bacteria were complement-opsonized (Fig. 1B-C; Suppl. Video 2-3). Most kidney MNPs express CD11b and CD11c, which are components of complement receptors CR3 and CR4, respectively, which mediate uptake of opsonized material. To determine whether these receptors mediate uptake of opsonized UPEC, we blocked them with antibodies. Blocking either receptor individually partially reduced bacterial phagocytosis, and blocking both of them was most effective (Fig. 1E).

**Fig 1.**
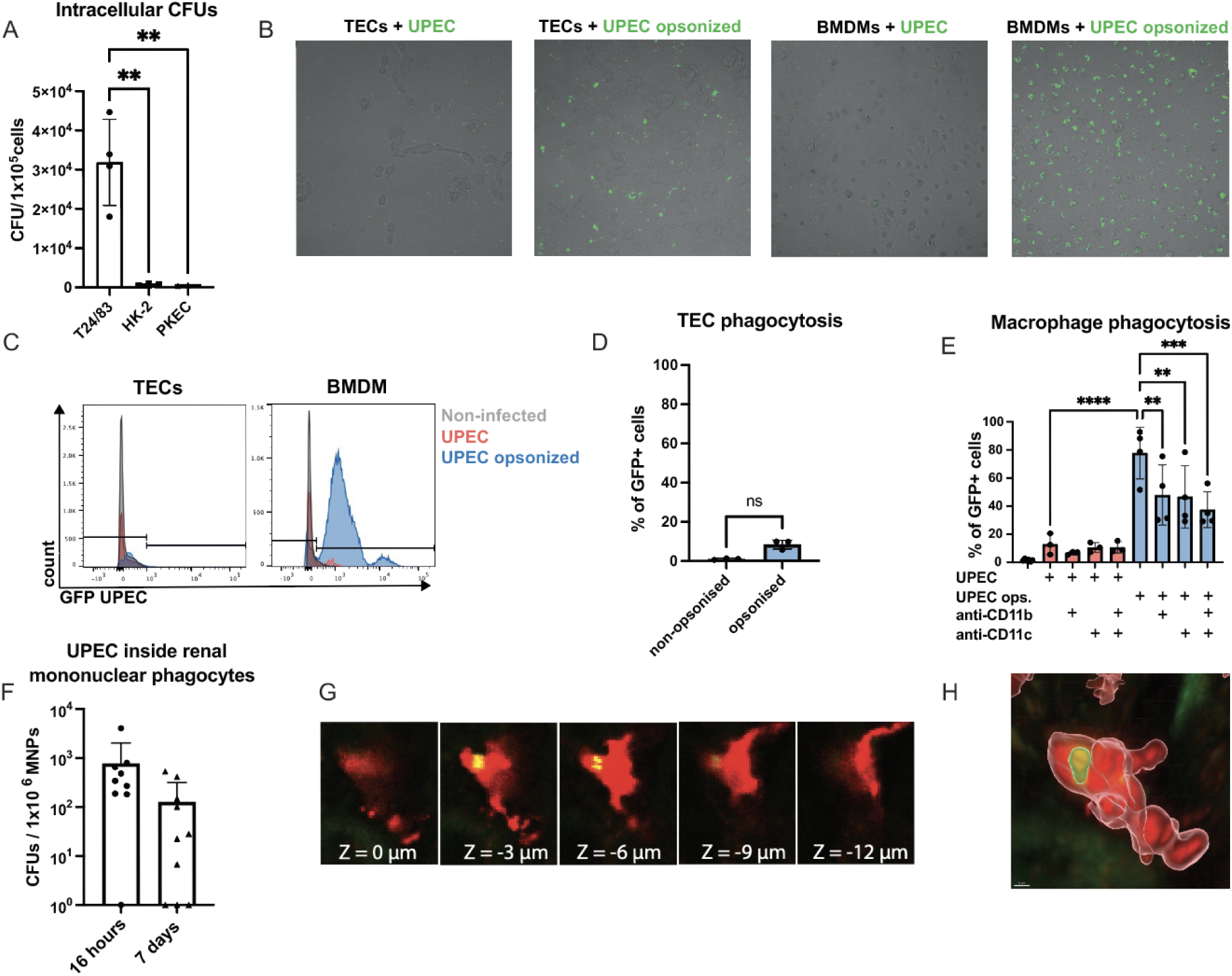
UPEC enter mononuclear phagocytes and bladder uroepithelial cells much better than kidney tubular epithelial cells. (A) CFU of bladder and kidney intracellular UPEC. HK-2 kidney proximal tube epithelial cells, and primary kidney epithelial cells (PKEC) were infected as with UPEC CFT073 for 1h at MOI of 50 and treated with gentamycin for 3 h. n = 3-4, 2 independent experiments. (B) Live cell imaging of opsonized and non-opsonized UPEC (green) cultured with tubular epithelial cells (TECs) and bone marrow-derived macrophages (BMDMs). (C) Representative histograms of opsonized (blue) and non-opsonized (red) UPEC phagocytosis by TECs and BMDMs. (D-E) Quantification of UPEC phagocytosis by TECs (D) and BMDMs (E) calculated as % of GFP+ cells. n = 3-4, 2 independent experiments. (F) CFU in renal mononuclear phagocytes (MNP) normalized to 1x10^6^ cells. n = 9-10, 3 independent experiments. (G) Intra-macrophage localization of UPEC. Vibratome-cut renal tissue section shows UPEC inside MNP after injecting UPEC into the ureter. Bright green, *E. coli* GFP (appears yellow inside MNPs); red macrophage, labeled with F4/80-PE. (H) 3D reconstruction of intramacrophage UPEC localization from (G). Bars show mean values; error bars indicate SD. \**P* < 0.05, \*\**P* < 0.01, and \*\*\**P* < 0.001 analyzed by One-way ANOVA (A), Mann-Whitney Test (D) and Two-way ANOVA (E).

These findings suggested that UPEC used CR3 and CR4 to establish intracellular colonies within renal MNPs, rather than in epithelial cells. We tested this hypothesis using our murine pyelonephritis model^28^ and isolated MNPs from mouse kidneys at 16 hours and 7 days post-infection, representing the early and advanced infection. Isolated MNPs were plated on LB agar, and bacterial colony-forming units (CFU) were enumerated. At 16 h post-infection, viable UPEC were present in MNPs from the kidneys of 8 of 9 mice (89%), with an average CFU of 776 (Fig. 1F). After 7 days, bacteria were still detected in the kidneys of 7 out of 10 mice (70%) with an average CFU of 126 (Fig. 1F). This was not limited to the UPEC strain 536, as also UPEC CTF073 entered and survived in MNPs (Fig. S1B). We visualized UPEC associated with renal MNPs using confocal imaging (Fig. 1G) and performed 3D-imaging reconstruction to highlight the intracellular localization of UPEC within renal MNPs from pyelonephritic mice (Fig. 1H). These findings confirmed that renal MNP can provide a niche for UPEC *in vivo*.

### Reduced invasion of itgam/itgax^−/−^ macrophages by UPEC

We next asked whether UPEC used CR3 and CR4 to enter macrophages also *in vivo*. The observation that they used both receptors to enter MNP *in vitro* (Fig. 1E) implied that, for *in vivo* loss-of-function experiments, CR3 and CR4 receptors need to be deleted. CD18 knockout mice cannot be used for this purpose, because they lack not only the integrin partner of CD11b and CD11c, but also of CD11a, which profoundly alters the architecture of lymphoid tissues and the immune cell composition^32^. Crossing CD11b and CD11c knockout mice was not feasible either, because these genes are adjacent on the same chromosome. Therefore, we used CRISPR/CAS to remove the entire CD11b/CD11c locus (Fig. S2A, B).

We first characterized the immune cell composition of *Itgam/Itgax^−/−^*mice by flow cytometry. A challenge was the absence of CD11b and CD11c, which are widely used MNP subset markers. Instead, we used CD172 (SIRP1a) as a surrogate for CD11b and the absence of CD64 for CD11c expression. In combination with MHC II, this revealed that *Itgam/Itgax^−/−^* mouse line showed no major alterations of myeloid immune cells, including neutrophils, and of lymphoid cells in kidney, spleen and lymph nodes (Fig. S3D-F). The architecture of their spleens and lymph nodes was comparable to WT mice (Fig. S3A, B).

Subsequently, we compared bacterial uptake by WT and *Itgam/Itgax^−/−^*bone-marrow-derived macrophages by exposing them to opsonized UPEC and performing live cell imaging. After 3 hours, nearly all WT macrophages were associated with UPEC, whereas only half of the knock-out macrophages showed a similar association (Fig. 2A-C; Suppl. Video 4, 5). Additionally, free UPEC in the supernatant of *Itgam/Itgax^−/−^* macrophages tended to be higher than from WT macrophages (Fig. 2D). When these macrophages were co-cultured overnight with gentamicin to kill the extracellular UPEC, more intracellular viable UPEC could be recovered from WT than from *Itgam/Itgax^−/−^* mice (Fig. 2E). These findings confirmed *in vitro* that *Itgam* and *Itgax* act as entry receptor for complement-opsonized UPEC.

**Fig 2.**
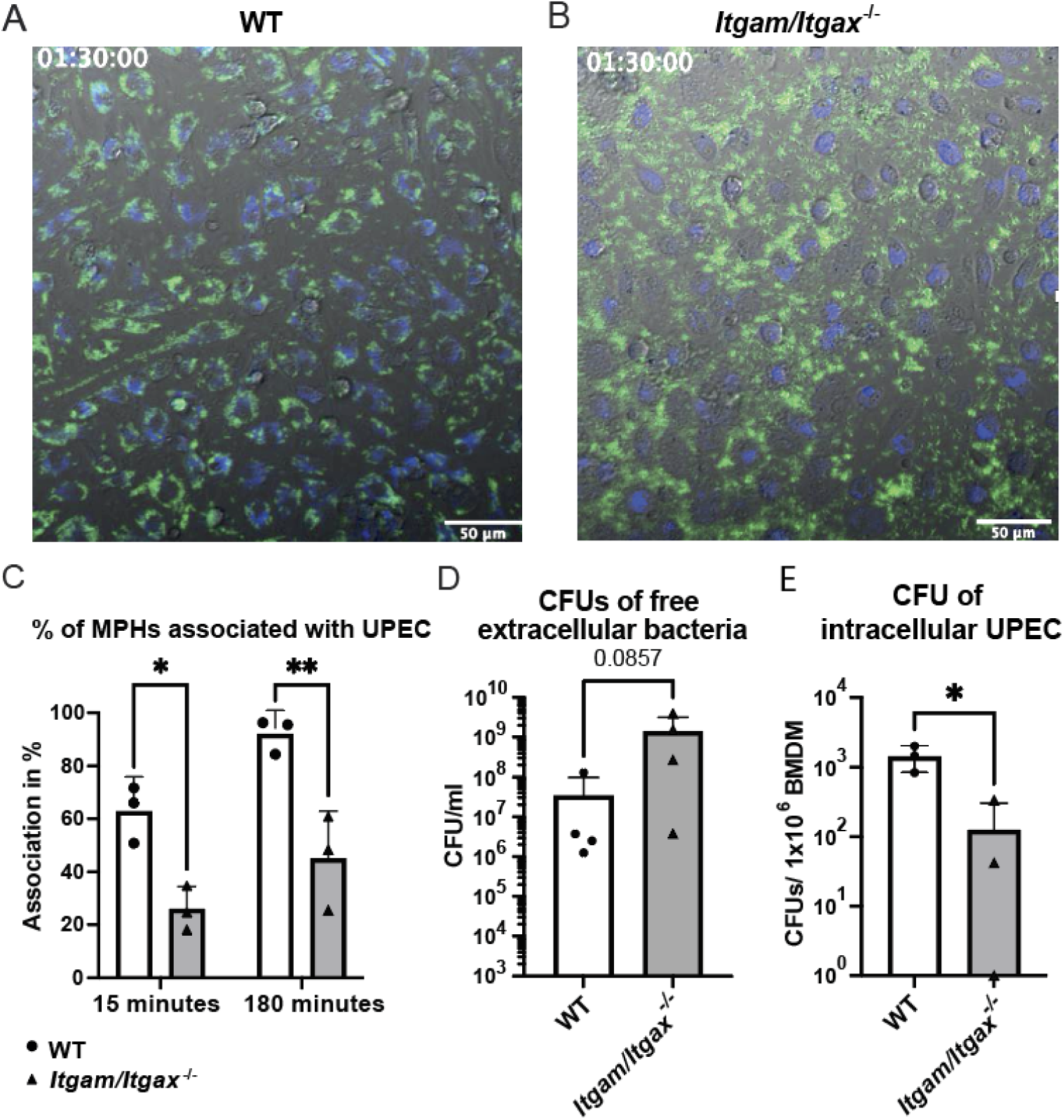
*Itgam* and *Itgax* are necessary for establishment of macrophage-UPEC associations. (A, B) *Ex vivo* live cell imaging of bone marrow-derived macrophages (BMDMs) (blue; F4/80-AF647) isolated from WT and *Itgam/Itgax*-deficient macrophages co-cultured with plasma-opsonized UPEC (green). (C) Associations of WT and *Itgam/Itgax*-deficient macrophages with UPEC at different imaging time points (15 and 180 min). n = 3, 3 independent experiments. (D) CFUs in the supernatant post-imaging. n = 4, 4 independent experiments (E) UPEC viability within mononuclear phagocytes determined by CFU enumeration. n = 3, 3 independent experiments. Extracellular bacteria were killed with gentamicin. Bars show mean values; error bars indicate SD. \**P* < 0.05, \*\**P* < 0.01, and \*\*\**P* < 0.001 analyzed by Two-way ANOVA (C), Mann-Whitney Test (D) and Unpaired t test (E).

### *Itgam/Itgax*^−/−^ mice clear bacterial pyelonephritis better than WT mice

To investigate the roles of CD11b and CD11c in pyelonephritis, we infected *Itgam/Itgax^−/−^* and WT mice and analyzed their kidneys. After 7 days, the renal bacterial load was reduced 100-fold in *Itgam/Itgax^−/−^*mice (Fig. 3A). Numbers of macrophages and neutrophils were unaltered (Fig. 3B-C), only numbers of cDC1s and cDC2s (conventional dendritic cells 1 and 2) in *Itgam/Itgax^−/−^* mice were lower at this time point, possibly reflecting their departure to the draining lymph node during infection (Fig. 3D, E). Expression of TNF-α, IL-6 and IL-1β was lower in the knock-out group, indicating reduced inflammation (Fig. 3F-H). When we analyzed F4/80^+^ MNP isolated from infected *Itgam/Itgax^−/−^* mice for intracellular UPEC survival, hardly any CFU could be detected, in contrast to WT controls (Fig. 3I), confirming that UPEC use CD11b and CD11c to enter and reside within renal MNP *in vivo*.

**Fig. 3.**
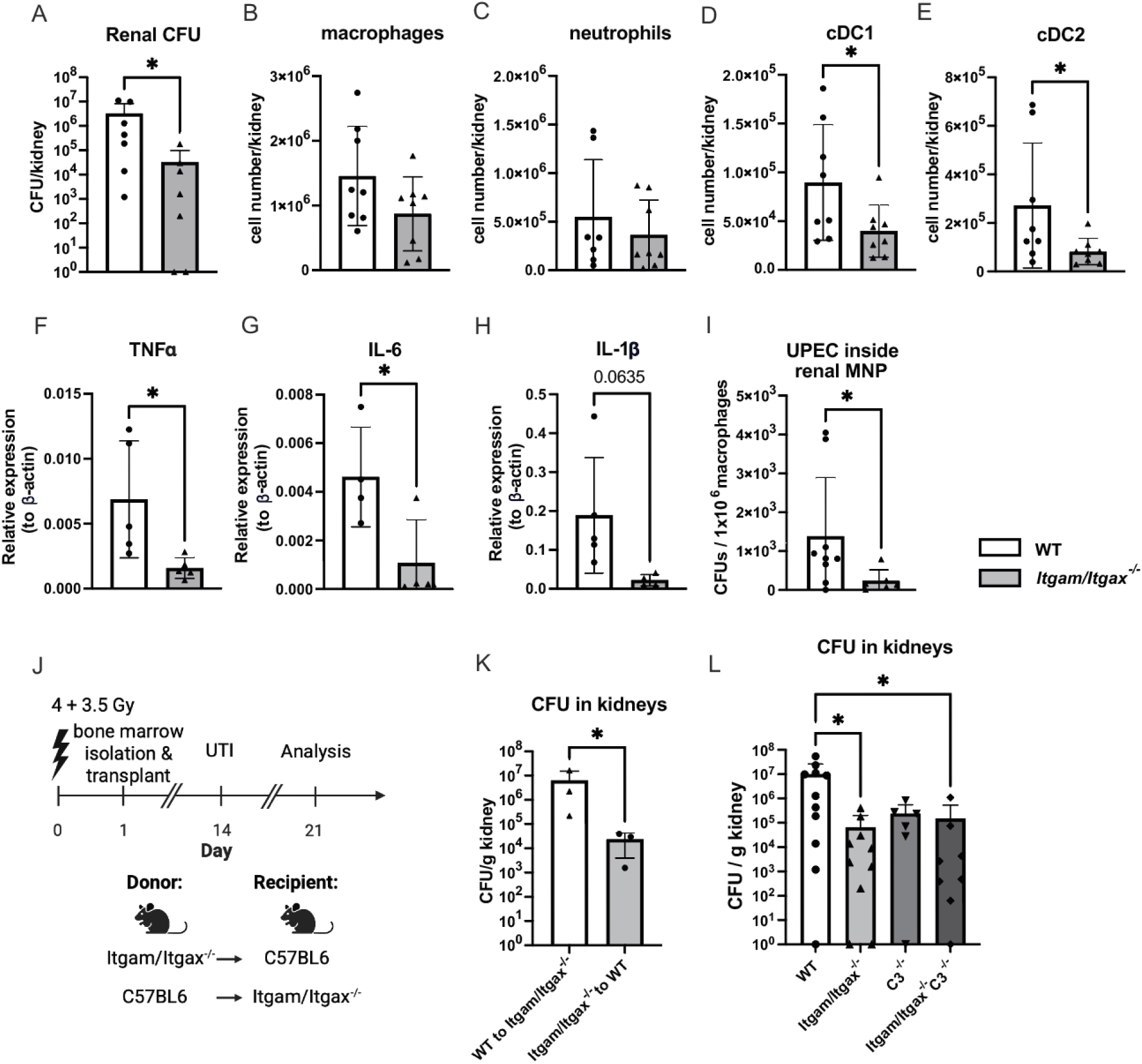
Pyelonephritis is attenuated in *Itgam/Itgax^−/−^* mice. (A) Bacterial load in kidneys at day 7 post-infection, (B-E) Intrarenal immune cell populations after 7 days measured by flow cytometry. (F-H) Relative expression of inflammatory cytokines in kidneys measured by RT-qPCR and represented as ∂Ct values. (I) Quantification of *E. coli* colony-forming units (CFU) in renal mononuclear phagocytes (RM). (J) Experimental timeline of UTI in bone marrow chimeras. (K) Bacterial burden in the kidneys of *Itgam/Itgax^-/-^* mice reconstituted with C57BL/6 bone marrow and C57BL/6 mice reconstituted with *Itgam/Itgax^-/-^*bone marrow. n = 3, one independent experiment. (L) Bacterial burden in kidneys of WT, *Itgam/Itgax^−/−^, C3^−/−^*and *Itgam/Itgax^−/−^x C3^−/−^*mice. 7 days post-infection, kidneys were harvested, and F4/80 positive isolation was performed to extract RM, which were subsequently plated on LB agar. n = 6-7, two independent experiments. Bars show mean values; error bars indicate SD. \**P* < 0.05, \*\**P* < 0.01, and \*\*\**P* < 0.001 analyzed by t test (B-E), Unpaired t test (F-H), Mann-Whitney Test (A, I, K), Kruskal-Wallis test (L).

To document that only CD11b^+^ CD11c^+^ immune cells constituted the intrarenal cellular niche, we transplanted bone marrow from WT mice to *Itgam/Itgax^-/-^* recipients and *vice versa* (Fig. 3J). Renal MNP from infected *Itgam/Itgax^-/-^* mice reconstituted with WT bone marrow showed high CFU numbers, but not MNP from WT mice reconstituted with *Itgam/Itgax^-/-^* bone marrow (Fig. 3K), arguing against non-hematopoietic cells as an additional intrarenal niche for UPEC.

To demonstrate that the reduced bacterial load in kidneys of *Itgam/Itgax^−/−^*mice was due to complement-opsonized bacteria binding to CD11b and CD11c, we crossed *Itgam/Itgax^−/−^* mice with mice deficient for the central complement component C3. The reduction of the renal bacterial load in *Itgam/Itgax^−/−^*x C3*^−/−^* mice was comparable to that in *Itgam/Itgax^−/−^* and in C3*^−/−^*mice (Fig. 3L), confirming that opsonized bacteria mainly use CR3 and CR4 to enter MNP and that this entry is the dominant mechanism of complement-mediated aggravation of pyelonephritis.

### Blocking CD11b and CD11c improves antibiotic therapy

We hypothesized that the MNPs niche might allow UPEC to escape antibiotics that cannot penetrate cell membranes, such as beta-lactams like Cephalosporins that are presently considered first line regimens in pyelonephritis^16,17^. To investigate this hypothesis, we treated infected WT mice every 12h starting at 24h post-infection, until 7 days after infection with Cefpodoxime, a 3^rd^ generation Cephalosporin and, as a control, with Ciprofloxacin that can enter cells, and analyzed kidneys 14 days after infection (Fig. 4A). Indeed, Cefpodoxime failed to eliminate UPEC from the kidney, whereas Ciprofloxacin treatment did so in all 8 mice (Fig. 4B). By contrast, cefpodoxime was able to significantly reduce the mean renal CFU number in *Itgam/Itgax^−/−^*mice (Fig. 4C), indicating that the intracellular MNP niche provides refuge from non-membrane penetrable antibiotics. If so, then combining cefpodoxime treatment with CD11b and CD11c blockade to prevent MNP re-entry should have a similar effect in WT mice with pyelonephritis. We first confirmed that antibodies against these integrins were effective in the kidney for 4 and 7 days (Fig. 4D-E). In infected mice, such blockade reduced the infection rate from 67% to 45% of mice, and the bacterial burden was similar as in the controls (Fig. 4G). Importantly, combining the blocking antibodies with the extracellularly acting antibiotic significantly decreased the infection rate to 1 of 11 mice, and the bacterial burden as well (Fig. 4G). In addition, renal inflammation after 2 weeks was attenuated as manifested by reduced numbers of immune cells (Fig. 4 H-I). CFUs in the bladder did not change after any treatment (Fig. 4J), which is consistent with previous findings showing that UPEC use uroepithelial cells to hide in this organ and use uroplakin, but not CD11b/c to enter^19^.

**Fig. 4.**
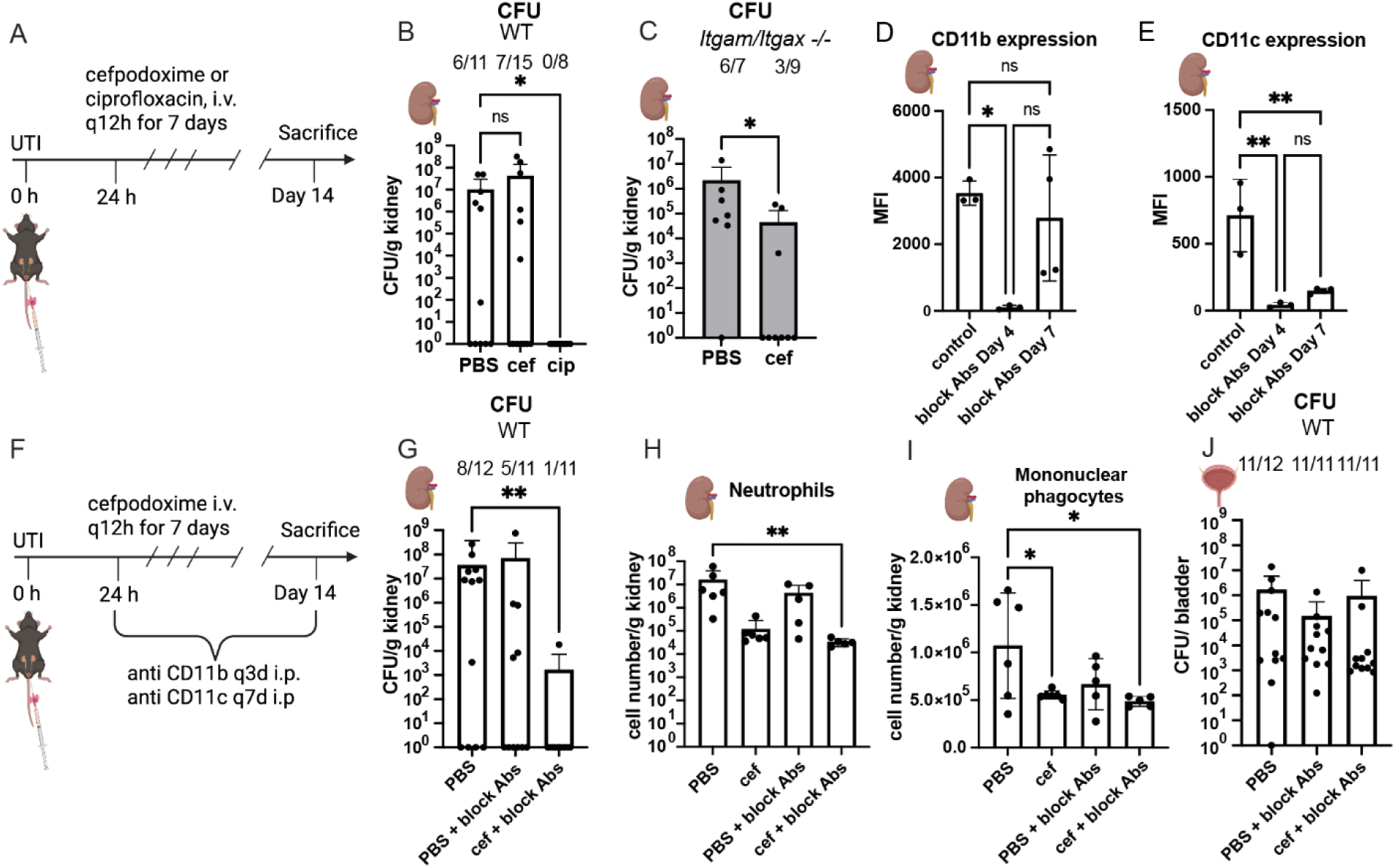
Blocking CD11b and CD11c improves antibiotic therapy in PN. (A) Experimental timeline for B and C. Mice infected with UPEC were treated with an extra-(cefpodoxime – cef) or intracellular (ciprofloxacin – cip) acting antibiotic. Antibiotics were administered *i.v.* for 7 consecutive days. (B, C) Bacterial load in the kidneys of WT (B) and *Itgam/Itgax^−/−^*(C) mice 14 days post-infection after applying the intracellular antibiotic iprofloxacin or the extracellular antibiotic cefpodoxime. n = 7-15, 3 independent experiments. (D-E) Mice were treated with blocking antibodies (anti-CD11b and anti-CD11c) *i.p*. CD11b remained blocked in the kidney for up to 4 days (D), whereas CD11c blockage persisted for more than 7 days (E). (F) Experimental timeline for G and H. Mice infected with UPEC were treated with cefpodoxime for 7 consecutive days, together with the blocking antibodies. Bacterial load in the kidneys (G) and bladder (F) 14 days post-infection. Treatment with extracellular antibiotics in combination with blocking antibodies protects mice from chronic pyelonephritis. n = 11-12, 2 independent experiments. Numbers above the bars indicate how many animals were used for the experiments and how many of them were still infected. The numbers of neutrophils (I) and mononuclear phagocytes (J) quantified by flow cytometry in untreated mice, mice treated with cefpodoxime, mice receiving blocking anti-bodies and mice receiving both treatments. n = 5-6, 2 independent experiments. Bars show mean values; error bars indicate SD. \**P* < 0.05, \*\**P* < 0.01, and \*\*\**P* < 0.001 analyzed by Kruskal-Wallis test (B, G, H), Mann-Whitney test (C), Ordinary one-way ANOVA (D, E, J) and Kruskal-Wallis test (I). Bar graph indicate means.

## DISCUSSION

Intracellular bacterial reservoirs are thought to contribute to relapse of various infections by shielding pathogens from immune effectors and many antibiotics. In the urinary tract, intracellular colonies of UPEC have been described within bladder uroepithelial cells and proposed to promote recurrent cystitis^32^. UPEC also form intracellular colonies in the kidney, but attempts to identify the cellular niche have been inconclusive so far. Here we identify renal MNPs as the principal intracellular reservoir in the kidney for UPEC *in vivo* and identify complement receptor-mediated uptake as entry pathway enabling this niche. Complement opsonization of UPEC has been implicated in their intrarenal persistence, but the underlying cellular mechanism remain unresolved, given that renal epithelial cells generally lack complement opsonin receptors. Our findings resolve this conundrum by demonstrating that complement-opsonized bacteria exploit receptors CR3 and CR4 on renal MNPs to gain intracellular access. Genetic ablation of the corresponding integrins Itgam and Itgax abolished intracellular bacterial persistence, establishing complement receptor–mediated uptake as a key host pathway enabling bacterial survival within the kidney. Unlike Itgb2 deficiency, which disrupts several leukocyte integrins and causes lymphoid abnormalities, selective deletion of Itgam and Itgax preserved immune architecture and allowed elucidating complement receptor function during pyelonephritis.

The exploitation of phagocytes as intracellular niches is a common strategy among bacterial pathogens. Our data indicate that UPEC employ that strategy in the kidney to evade neutrophils, the main immune effectors during pyelonephritis. Furthermore, sequestration within MNPs protects bacteria from antibiotics with limited cellular penetration, especially from β-lactams that remain first-line therapy for urinary tract infections. The relatively rapid turnover of renal MNPs imposes a dynamic aspect to this intracellular niche. In contrast to uroepithelial cells, which may harbor bacteria for prolonged periods, tissue macrophages undergo physiological replacement and are even more rapidly replenished during inflammation by recruited monocytes^26^. Consequently, intracellular bacteria must periodically transition from dying host MNP into new ones to maintain persistence. Our findings suggest that this transition represents a vulnerability that can be therapeutically exploited. Pharmacological inhibition of complement receptors prevented bacterial re-entry into MNPs and markedly enhanced the efficacy of extracellularly acting antibiotics. This observation may explain the necessity of prolonged treatment courses for β-lactam therapy in pyelonephritis, despite the rapid bactericidal activity of these drugs in vitro^33,34^.

Our findings suggest complement receptor blockade as the first host-directed strategy to potentiate antibiotic therapy in pyelonephritis. Such an approach may be particularly valuable in the context of the globally rising antimicrobial resistance, where therapeutic options become increasingly narrow. Other uropathogens, including *Klebsiella* species, possess similar outer membrane structures and are likewise susceptible to complement opsonization^35^, raising the possibility that complement receptor-mediated uptake may represent a broader mechanism exploited by Gram-negative bacteria during infection.

Several complement inhibitors are already being explored therapeutically in other inflammatory diseases^36^, but none of them selectively targets CR3 and CR4. Our findings suggest that the currently employed inhibitors, which target downstream complement components, such as C5 or anaphylatoxin receptors, are unlikely to prevent the establishment of intracellular bacterial reservoirs, because they prevent neither C3b-mediated opsonization nor receptor engagement. Blocking CR3 and CR4 may provide a more targeted approach that preserves other complement effector functions, such as anaphylatoxin production essential for neutrophil recruitment, or membrane-attack complex formation to directly lyse bacteria, thereby limiting immunosuppressive side-effects.

Taken together, our findings identify the inhibition of complement receptor-mediated uptake into MNP as a novel therapeutic option to prevent intracellular bacterial persistence during pyelonephritis. This strategy exposes bacteria to immune effector cells and extracellular antibiotics, and therefore may represent a promising adjunct strategy to improve treatment of recurrent urinary tract infections in the era of increasing antimicrobial resistance. This principle may extend to infections by further bacteria known to exploit complement receptors^37^.

## Disclosure

All the authors declared no competing interests.

## Data statement

All data are available from the lead contact on request.

## Acknowledgments

We thank V. Kotov and M. Blankert for technical assistance. We acknowledge the support by the Central Animal Facility and the Flow Cytometry Core Facility of the Medical Faculty at Bonn University

## Funding

This project was supported by funding from the German Research Foundation FOR 5427/1 ‘‘BARICADE’ (project number 466687329) to FW, UD, DRE, CK, SJ and SvV and by a Gottfried Wilhelm Leibniz-Award to CK. CK, SvV and NG are members of the Excellence Cluster ImunoSensation³ EXC2151 project number 390873048.

## Author contributions

Conceptualization: CK, SKJ, BG, FW; Methodology: MH, CK, SKJ, FW, SvV, DRE, NG, UD; Investigation: BG, OB, PR, YM, MS, CL, NB, DK, AV, AA, JY; Visualization: BG, OB, PR; Funding acquisition: CK, SKJ, FW; Supervision: CK, SKJ, FW; Writing – original draft: BG, SKJ, CK, FW; Writing – review & editing: all authors

## Competing interests

Authors declare that they have no competing interests.

## Data and materials availability

All data are available in the main text or the supplementary materials.

## MATERIAL AND METHODS

### Study design

Mouse experiments were conducted in compliance with Animal Care and Use Committee guidelines and had been approved by the relevant governmental authorities. Our study examined female mice because UTI occurs far more frequently in females. We defined the endpoint of the *in vivo* experiments at 2- and 7- and 14 days post-infection. Following infection, only mice with positive for UPEC in urine at 12 hours post-infection were further included. Sample sizes for each group were determined based on experience from previous or preliminary experiments and were subsequently confirmed through statistical power analysis. No data was excluded. Sample sizes are indicated in the figures or their legends and correspond to the number of animals. Figure legends indicate the performed statistical tests. All results show the mean of at least 3 independent samples per group.

### Animals

8- to 12-week-old C57BL/6 mice were bred in-house and accommodated in a specific pathogen-free unit at the Institute of Molecular Medicine and Experimental Immunology in Bonn. All mice had access to autoclaved tap water and palleted food ad libitum. Animal experiments had been approved by a local animal ethics reviewing board. ARRIVE guidelines were followed.

### Generation of *ItgamItgax*^-/-^ mouse line

The *ItgamItgax*^-/-^ mouse line was created in cooperation with Prof. Marco Herold in the core facility MAGEC of the WEHI in Melbourne, Australia. The mouse line was generated using CRISPR/Cas9 technology. To delete sequences of *Itgam* and *Itgax* genes (located on chromosome 7 side by side), two sgRNAs with the following sequences were designed: TAACCCTGATGGTTCGGGCC and CTCCCAATAGTTTCGGTATC. For deletion was targeted 89, 984 bp of genomic sequence. First, Cas9 and both sgRNAs were injected into single embryos with a C57BL/6 background. Then, these embryos were transferred into female mice. After the birth, pups were checked for the presence of the deleted allele. For the detection of the deleted allele of F0 mice following primers were used: 5’-ACCAGCCTGATCCGAAACAC-3’ (sense) and 5’-GTGCTGCTTGTGGGAACTTG-3’ (antisense). PCR conditions were:

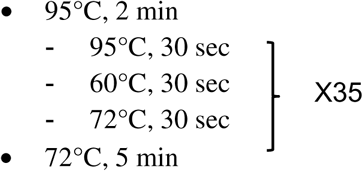

To detect the WT sequence, the following primers were used: 5’-AGGGAGAGGTAGGAAGAGGC-3’ (sense) and 5’-CTGCACCAGTGAGAGAGAGC-3’ (antisense). PCR was performed under the conditions listed below:

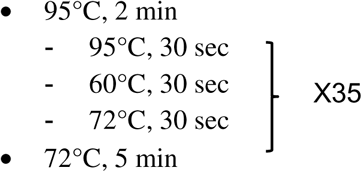

Material from the pups with a confirmed knockout of *Itgam* and *Itgax* was then used for next-generation sequencing to validate the deletion of the genes of interest. Mice in which deletions were confirmed were backcrossed with WT animals for 2 generations to breed out potential off-target mutations. Afterward, offspring were intercrossed and further bred to homozygosity

### Bacteria culture

Uropathogenic E. coli 536 stain with and without GFP, and CFT073 were provided by Ulrich Dobrindt from the University of Münster, Germany. Bacteria were grown from a single colony picked from a freshly streaked lysogeny broth (LB) agar plate and placed in 4 ml LB at 37°C with shaking (180 rpm) overnight. The following day, bacteria were subcultured in 100 ml LB until OD_600_ = 1,1 -1,2, centrifuged at 4000 rpm for 20 min and resuspended in saline at 10^10^ CFU in 1 mL before transurethral injection.

### Opsonization of UPEC

To opsonize *UPEC-GFP* strain 536, an overnight culture was prepared by inoculating the bacteria in 3 mL of LB medium and incubating at 37°C with shaking at 180 rpm O/N.

On the day of the experiment, the overnight culture was transferred to a 15 mL conical tube and centrifuged at 3,000 rpm for 5 minutes at 4°C. The bacterial pellet was resuspended in 1 mL of sterile PBS, and the concentration was determined using Trypan blue exclusion (1:200 dilution) with a Neubauer chamber. The bacterial suspension was then adjusted to a MOI of 10 (1.2 × 10□ bacteria per 20 μL of macrophages) using PBS in a 50 mL tube.

Following adjustment, 500 μL of the bacterial suspension was aliquoted into two 1.5 mL tubes and centrifuged at 12,000 rpm for 5 minutes at 4°C. For opsonization, each pellet was resuspended in 500 μL of PBS++ (1.2 mM CaCl□ and 4.25 mM MgCl□) supplemented with 100 μL of 20% murine serum. For the non-opsonized control, 600 μL of PBS was added to the pellet and resuspended. All tubes were incubated at 37°C for 30 minutes with shaking at 180 rpm.

### Live-cell imaging of TEC and bone marrow-derived macrophages

To facilitate visualization of tubular epithelial cells (TEC) and bone marrow-derived macrophages (BMDM) using the Zeiss LSM710 inverted confocal laser scanning microscope, cells were plated in an 8-well ibidi chamber one day prior to imaging. Macrophages were harvested on day 6 of culture. A staining master mix was prepared consisting of F4/80-AF647 (1:300), Fc block (1:33), and FACS buffer, calculated at 50 μL of master mix per 1 × 10□ cells. The cells were seeded into the ibidi chamber at a density of 1.2 × 10□ cells per 100 μL per well, according to the imaging setup and infected with UPEC 536 MOI of 10 opsonized with plasma.

To facilitate macrophage attachment and growth, the chamber was centrifuged at 300×g for 1 minute at 4°C, then placed in an incubator overnight at 37°C with 5% CO□. HK-2 cell line was cultured in DMEM/F12 medium. 1 × 10□ cells per 100 μL per well were seeded into an 8-well ibidi chamber and infected with UPEC 536 MOI of 10 opsonized with plasma.

### Mouse infections

8-to 12-week-old female mice were anaesthetized with ketamine (200 mg/kg) and xylazine (10 mg/kg), placed on a heating pad and infected by transurethral instillation of 10^9^ UPEC with a flexible polyethylene catheter (outer diameter 0,6 mm, DB) covered with Instillagen (Farco Pharma). The infection was repeated after 3 hours to induce pyelonephritis.

### Bacteriological analysis of murine kidneys and bladders

The number of bacteria in mice was quantified by scoring CFU after kidneys and bladders homogenization in PBS with an Ultra Turrax. For determination of CFU, tissue homogenate was serially diluted and plated in a volume of 20 μL on LB agar. Agar plates were incubated at 37°C O/N. The bacterial colonies were counted the next day.

### Immune cells isolation

Mice were euthanized in a CO_2_ chamber. Harvested kidneys were placed in a 24-well plate, with each well containing 1 mL of digestion medium (RPMI medium supplemented with 1% DNase and 1% collagenase IV) and homogenized using a 2 mL syringe plunger. The homogenates were incubated at 37°C for 1 hour, then resuspended with a 1 mL pipette. The resulting solutions were filtered using a 100 μL filter, rinsed with 1 mL of FACS buffer, and transferred into 15 mL Falcon tubes. The samples were centrifuged at 300 rpm for 2 minutes, and the supernatants were transferred to new 15 mL Falcon tubes, which were filled to 5 mL with FACS buffer. The, the solutions were spun down at 1300 rpm for 5 minutes, and supernatants were removed. Samples were resuspended in 1 mL of RBC lysis buffer, and the reaction was stopped by adding FACS buffer to 10 mL. Tubes were centrifuged at 1300 rpm for 5 minutes, and supernatants were removed. Finally, cells were resuspended in 500 μL of FACS buffer and used for surface or intracellular staining.

### Flow Cytometry

BD LSRFortessa, housed in the Flow Cytometry Core Facility at the Medical Faculty of University of Bonn, was used for this study. This cytometer consists of five lasers (355nm, 405nm, 488nm, 561nm and 640nm) and allows for the simultaneous analysis of up to 20 parameters. Sample acquisition was performed in BD FACSDiva software v9.0 and the data were analyzed in FlowJo 10.8.2. Antibodies and reagents used for flow cytometry are summarized in table below.

| Antigen | Reactivity | Host | Fluoro chrome | Clone | Isotype | Company |
| --- | --- | --- | --- | --- | --- | --- |
| CD103 | mouse | hamster | APC | 2E7 | Armenian Hamster IgG | BioLegend |
| CD11b | mouse | rat | BUV395 | M1/70 | Rat IgG2b κ | Thermo Fisher |
| CD11c | mouse | hamster | BUV737 | HL3 | Armenian Hamster IgG1, λ2 | BD Bioscience |
| CD45 | mouse | rat | BV510 | 30-F11 | Rat IgG2bκ | BioLegend |
| CX3CR1 | mouse | mouse | BV421 | SA011F11 | Mouse IgG2aκ | BioLegend |
| F4/80 | mouse | rat | PerCP-Cy5.5 | BM8 | Rat IgG2a κ | BioLegend |
| Ly6C | mouse | rat | PE/Cy7 | HK1.4 | Rat IgG2c κ | BioLegend |
| Ly6G | mouse | rat | PE | 1A8 | Rat IgG2b κ | BioLegend |
| MHCII | mouse | rat | AF700 | MH/114.15.2 | Rat IgG2b κ | BioLegend |
| B220 | mouse | rat | PE | RA3-6B2 | Rat IgG2a κ | BioLegend |
| CD4 | mouse | rat | PerCp-Cy5.5 | GK1.5 | Rat IgG2b κ | BioLegend |
| CD8 | mouse | rat | APC | 53-6.7 | Rat IgG2a κ | BioLegend |
| CD8 | mouse | rat | BV510 | 53-6.7 | Rat IgG2a κ | BioLegend |
| CD19 | mouse | rat | APC-Cy7 | 6D5 | Rat IgG2a κ | BioLegend |
| CD172a | mouse | rat | PE/Dazzle5P4 | P84 | Rat IgG1, κ | BioLegend |

### Isolation of bone marrow cells

Bone marrow-derived macrophages were isolated using a centrifugation-based method. Mice were euthanized in a CO_2_ chamber. Blood was collected from the heart using a sterile 1 mL syringe and a needle. The skin near the hindlimbs was opened with sterile forceps and scissors, and the underlying muscle tissue was removed to expose the femur and tibia, which were then carefully excised. Residual muscle tissue was removed with a clean tissue, and the bones were placed in a petri dish containing 10 mL of ice-cold PBS with 1% FCS. Each bone was cut at the knee end and inserted facing downward into a 0.5 mL Eppendorf tube with a pre-punched hole. These tubes were placed inside 1.5 mL collection tubes containing 100 μL of PBS + 1% FCS. The assembly was centrifuged at 2,500×g for 1 minute at RT to collect the bone marrow into the lower tube. If marrow remained (indicated by red coloration), the opposite end of the bone was cut, and the centrifugation step was repeated until the bones appeared white. The collected marrow was resuspended in 1 mL of PBS + 1% FCS by pipetting up and down 10 times. The cell suspension was filtered through a 70 μm strainer (Greiner) into a 50 mL tube, followed by an additional rinse of the strainer with 2 mL PBS + 1% FCS. The filtered suspension was centrifuged at 300×g for 5 minutes at RT. Red blood cell lysis was performed by resuspending the pellet in 3–5 mL of ACK lysis buffer (depending on pellet size and color), incubating for 3 minutes, and then quenching with 20 mL of PBS + 1% FCS. The cells were centrifuged at 1,250 rpm for 7 minutes at 4°C, and the resulting pellet was resuspended in PBS (bone marrow transplantation) or 1 mL of DMEM (Gibco) supplemented with 20% M-CSF-containing supernatant from L929 cell culture (macrophage differentiation). Cell concentration was determined by Trypan blue exclusion (1:20 dilution) using a Neubauer chamber.

For macrophage differentiation, based on the cell counts, the suspension was distributed equally into 3–4 petri dishes, each containing 15 mL of DMEM + 20% M-CSF. The cells were incubated at 37°C with 5% CO□ for differentiation. On day 3, 5 mL of fresh DMEM + 20% M-CSF was added to each dish. Cell growth and contamination were monitored regularly.

On day 7, differentiated macrophages were harvested for downstream applications, including live-cell imaging and flow cytometry.

### Kidney imaging

To visualize macrophages that captured E. coli-GFP in the kidneys, mice were first anesthetized and injected with 10 μL of F4/80–AF647 and CD31–PE. Simultaneously, 50 μL of an overnight E. coli culture (OD: 1–1.1) was injected into the ureter of each kidney. After 30 minutes, the mice were euthanized via CO□ inhalation, followed by perfusion with 10 mL of saline and 5 mL of bath solution (150 mM NaCl, 5 mM KCl, 1 mM CaCl□, 2 mM MgCl□, 5 mM glucose, and 10 mM HEPES). Kidneys were harvested and placed in complete medium (phenol-red-free DMEM supplemented with 10% heat-inactivated FBS and 10 mM HEPES). Subsequently, kidneys were embedded in 4% low melting point agarose (Promega, V2111) until solidified. They were then sectioned into 200 μM slices using a vibrating microtome (Leica VT1000S; speed: 5, frequency: 7). Tissue sections were transferred to ibidi chambers and imaged using a 20x objective lens.

### Generation of bone marrow chimeras

C57BL/6 and *Itgam/Itgax^-/-^* mice were irradiated with two doses of gamma radiation (4 Gy and 3.5 Gy) at 4 hours interval, to ablate its bone marrow. The next day, they received an injection of donor bone marrow (2 million cells in 100 ul PBS *i.v.*) to establish a chimeric state.

### Statistics

Statistical differences were assessed using a two-tailed unpaired or paired Student’s t-test for comparisons between two groups. For comparisons involving three groups, a repeated-measures one-way ANOVA was applied. In the case of non-parametric data, the Mann-Whitney test was used for two-group comparisons, and the Kruskal-Wallis test for three or more groups. All statistical analyses were performed using GraphPad Prism 10. Statistical significance was defined as p < 0.05 (*).

## Supplementary Materials

### Supplementary data

**Fig S1.**
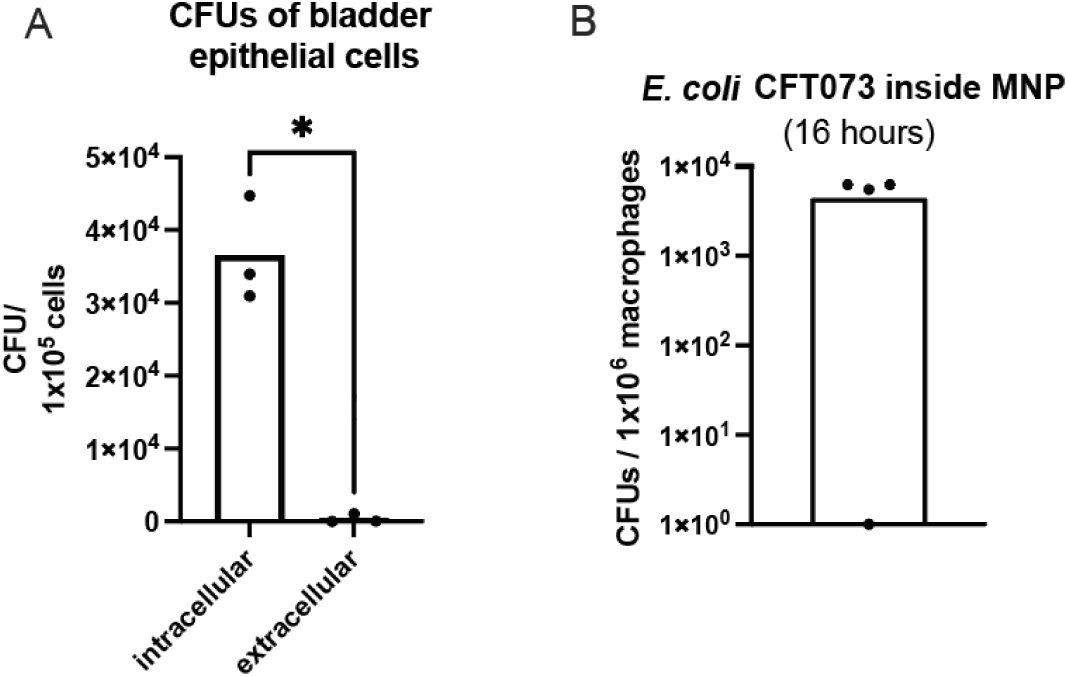
Intracellular survival of UPEC during experimental PN. (A) CFU of bladder intra- and extracellular UPEC. T24/83 bladder epithelial cell were infected with UPEC CFT073 for 1h at an MOI of 50 and treated with gentamycin for 3h. n = 3, 3 independent experiments (B) Mice were infected transurethrally with UPEC CFT073 to induce PN. Sixteen hours post-infection, kidneys were harvested and F4/80 positive isolation was performed to extract RM, which were subsequently plated on LB agar. n = 4, one independent experiment analyzed by Mann-Whitney test. Total CFUs were normalized to 1x10^6^ macrophages.

**Fig S2.**
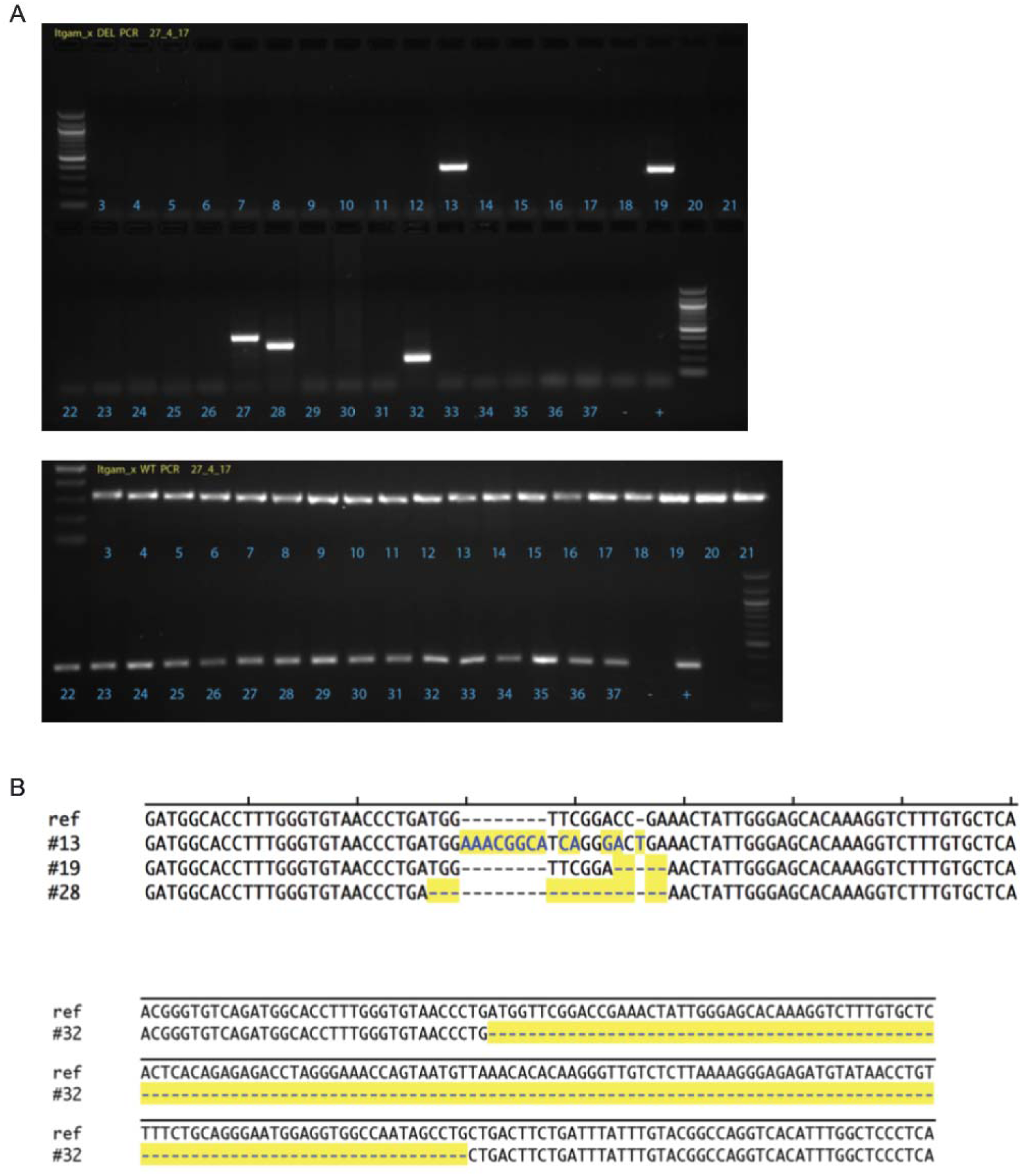
Generation of Itgam/Itgax^−/−^ mice. (A) Genotyping of F0 mice: A PCR product of 339 bp is expected if the deleted allele is present. A PCR product of 320 bp is expected if the wildtype allele i present. F0 mice #13, #19, #27, #28 and #32 were positive for the deleted allele and DNA samples from these mice were used for next generation sequencing to validate the deletions. (B) Next-generation sequencing of F0 offspring of mice positive for the deleted allele. Sequences of deleted alleles were aligned to a predicted deleted sequence. Following Cas9 cutting at the two sgRNA cut sites, the intervening genomic DNA sequence is excised and the early embryo often repairs this break with InDel mutations between the 2 sgRNA cut sites. These additional InDel mutations are highlighted in yellow.

**Fig S3.**
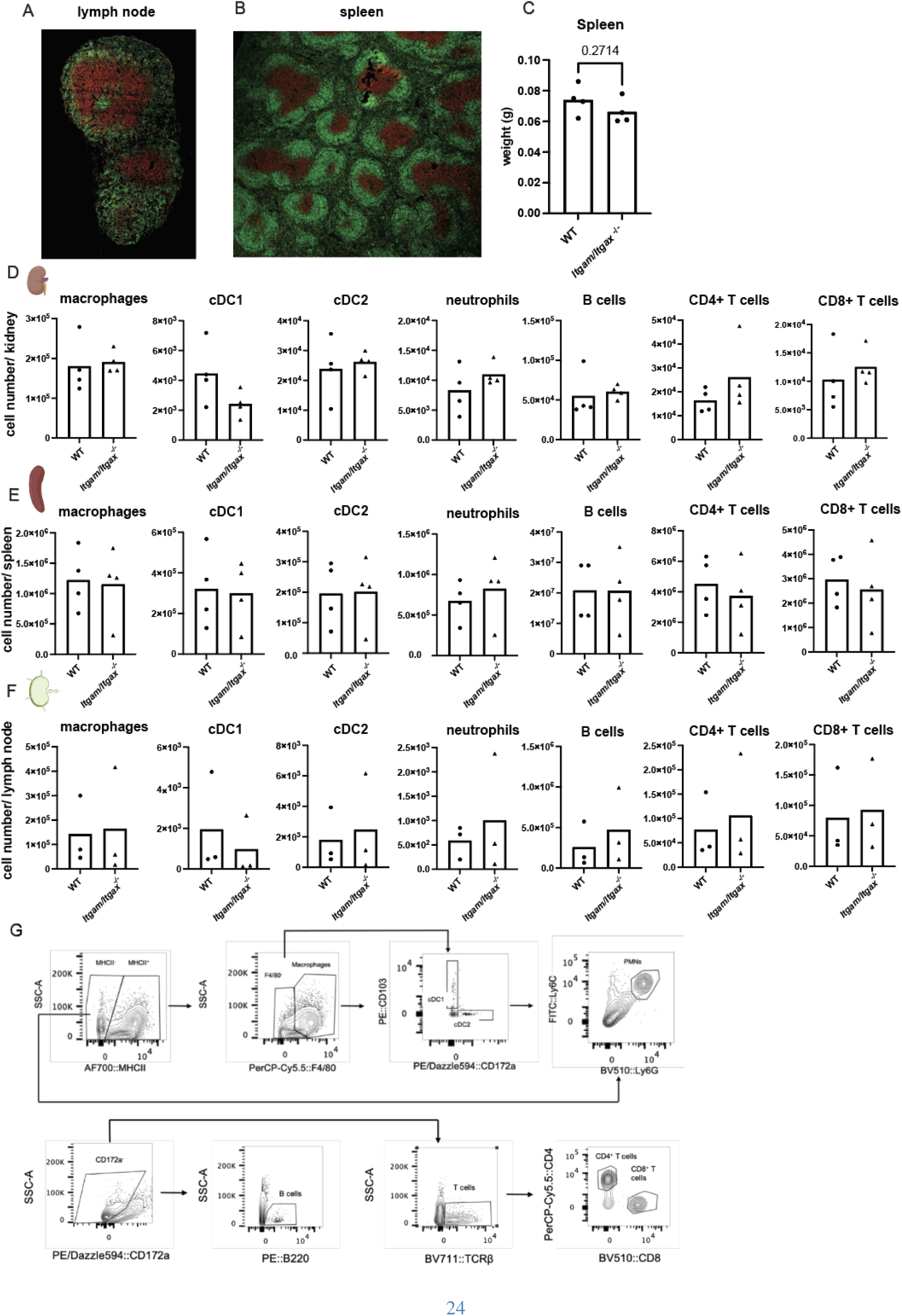
*Itgam/Itgax^−/−^*mice show no major alterations of immune cells in the kidney, spleen and lymph nodes. The architecture of lymph nodes (A) and spleen (B) is defined by distinct B cells (green, B220) and T cells (red, CD3) zones, typical for WT mice. (C) Spleen weight does not differ in WT and *Itgam/Itgax^−/−^* mouse. (D-F) No significant differences in myeloid or lymphoid immune cell populations were observed in the kidneys, spleens, or lymph nodes of *Itgam/Itgax^−/−^*mice. Immune cells were analyzed by flow cytometry. (G) Gating strategy for myeloid and lymphoid cells in the kidney and lymph nodes. \**P* < 0.05, \*\**P* < 0.01, and \*\*\**P* < 0.001 analyzed by unpaired t test. Bar graphs indicate means.

## Notes

### Competing Interest Statement

The authors have declared no competing interest.

## REFERENCES

1. Flores-Mireles AL, Walker JN, Caparon M, et al. Urinary tract infections: epidemiology, mechanisms of infection and treatment options. Nat Rev Microbiol. 2015;13(5):269–284.

2. Geerlings SE, Beerepoot MA, Prins JM. Prevention of recurrent urinary tract infections in women: antimicrobial and nonantimicrobial strategies. Infect Dis Clin North Am. 2014;28(1):135–147.

3. Graversen HV, Norgaard M, Nitsch D, et al. Preadmission kidney function and risk of acute kidney injury in patients hospitalized with acute pyelonephritis: A Danish population-based cohort study. PLoS One. 2021;16(3):e0247687.

4. Schwartz L, de Dios Ruiz-Rosado J, Stonebrook E, et al. Uropathogen and host responses in pyelonephritis. Nat Rev Nephrol. 2023;19(10):658–671.

5. Vermeulen SH, Hanum N, Grotenhuis AJ, et al. Recurrent urinary tract infection and risk of bladder cancer in the Nijmegen bladder cancer study. Br J Cancer. 2015;112(3):594–600.

6. Schreiber HLt, Conover MS, Chou WC, et al. Bacterial virulence phenotypes of Escherichia coli and host susceptibility determine risk for urinary tract infections. Sci Transl Med. 2017;9(382).

7. Antimicrobial Resistance C. Global burden of bacterial antimicrobial resistance in 2019: a systematic analysis. Lancet. 2022;399(10325):629–655.

8. McLellan LK, Hunstad DA. Urinary Tract Infection: Pathogenesis and Outlook. Trends Mol Med. 2016;22(11):946–957.

9. Mulvey MA, Lopez-Boado YS, Wilson CL, et al. Induction and evasion of host defenses by type 1-piliated uropathogenic Escherichia coli. Science. 1998;282(5393):1494–1497.

10. Dhakal BK, Kulesus RR, Mulvey MA. Mechanisms and consequences of bladder cell invasion by uropathogenic Escherichia coli. Eur J Clin Invest. 2008;38 Suppl 2:2–11.

11. Mysorekar IU, Hultgren SJ. Mechanisms of uropathogenic Escherichia coli persistence and eradication from the urinary tract. Proc Natl Acad Sci U S A. 2006;103(38):14170–14175.

12. Azimzadeh PN, Birchenough GM, Gualbuerto NC, et al. Mechanisms of uropathogenic E. coli mucosal association in the gastrointestinal tract. Sci Adv. 2025;11(5):eadp7066.

13. Flores C, Ling J, Loh A, et al. A human urothelial microtissue model reveals shared colonization and survival strategies between uropathogens and commensals. Science Advances. 2023;9(45):eadi9834.

14. Shrout JD. The Calming Effect of *Escherichia coli*. Science Translational Medicine. 2012;4(130):130ec166–130ec166.

15. Lehar SM, Pillow T, Xu M, et al. Novel antibody-antibiotic conjugate eliminates intracellular S. aureus. Nature. 2015;527(7578):323–328.

16. G. Bonkat JK, T. Cai, S.E. Geerlings, B. Köves, A. Pilatz, J. Medina-Polo, L. Schneidewind, S. Schubert, R. Veeratterapillay, F. Wagenlehner. Edn. presented at the EAU Annual Congress Madrid, Spain 2025. ISBN 978-94-92671-29-5. EAU Guidelines Office, Arnhem, the Netherlands. Edn. presented at the EAU Annual Congress Madrid, Spain 2025. 2025.

17. Barbara W. Trautner NWC-P, Kalpana Gupta, Elizabeth B. Hirsch, Molly Horstman, Gregory J. Moran, Richard Colgan, John C. O’Horo, Muhammad S. Ashraf, Shannon Connolly, Dimitri Drekonja, Larissa Grigoryan, Angela Huttner, Gweneth B. Lazenby, Lindsay Nicolle, Anthony Schaeffer, Sigal Yawetz, Valéry Lavergne. Clinical Practice Guideline by Infectious Diseases Society of America (IDSA): 2025 Guideline on Management and Treatment of Complicated Urinary Tract Infections: Selection of Antibiotic Therapy for Complicated UTI. 2025.

18. Ternes B, Wagenlehner FME. [Guideline-based treatment of urinary tract infections]. Urologe A. 2020;59(5):550–558.

19. Thumbikat P, Berry RE, Zhou G, et al. Bacteria-induced uroplakin signaling mediates bladder response to infection. PLoS Pathog. 2009;5(5):e1000415.

20. Schiwon M, Weisheit C, Franken L, et al. Crosstalk between sentinel and helper macrophages permits neutrophil migration into infected uroepithelium. Cell. 2014;156(3):456–468.

21. Uhlen M, Fagerberg L, Hallstrom BM, et al. Proteomics. Tissue-based map of the human proteome. Science. 2015;347(6220):1260419.

22. Thul PJ, Akesson L, Wiking M, et al. A subcellular map of the human proteome. Science. 2017;356(6340).

23. Karlsson M, Zhang C, Mear L, et al. A single-cell type transcriptomics map of human tissues. Sci Adv. 2021;7(31).

24. Springall T, Sheerin NS, Abe K, et al. Epithelial secretion of C3 promotes colonization of the upper urinary tract by Escherichia coli. Nat Med. 2001;7(7):801–806.

25. Aderem A, Underhill DM. Mechanisms of phagocytosis in macrophages. Annu Rev Immunol. 1999;17:593–623.

26. Ginhoux F, Schultze JL, Murray PJ, et al. New insights into the multidimensional concept of macrophage ontogeny, activation and function. Nat Immunol. 2016;17(1):34–40.

27. Liu F, Dai S, Feng D, et al. Distinct fate, dynamics and niches of renal macrophages of bone marrow or embryonic origins. Nat Commun. 2020;11(1):2280.

28. Tittel AP, Heuser C, Ohliger C, et al. Kidney dendritic cells induce innate immunity against bacterial pyelonephritis. J Am Soc Nephrol. 2011;22(8):1435–1441.

29. Nelson PJ, Rees AJ, Griffin MD, et al. The renal mononuclear phagocytic system. J Am Soc Nephrol. 2012;23(2):194–203.

30. Jobin K, Stumpf NE, Schwab S, et al. A high-salt diet compromises antibacterial neutrophil responses through hormonal perturbation. Sci Transl Med. 2020;12(536).

31. Godaly G, Ambite I, Puthia M, et al. Urinary Tract Infection Molecular Mechanisms and Clinical Translation. Pathogens. 2016;5(1).

32. Berlin-Rufenach C, Otto F, Mathies M, et al. Lymphocyte migration in lymphocyte function-associated antigen (LFA)-1-deficient mice. J Exp Med. 1999;189(9):1467–1478.

33. Cronberg S, Banke S, Bergman B, et al. Fewer bacterial relapses after oral treatment with norfloxacin than with ceftibuten in acute pyelonephritis initially treated with intravenous cefuroxime. Scand J Infect Dis. 2001;33(5):339–343.

34. Hyslop DL, Bischoff W. Loracarbef (LY163892) versus cefaclor and norfloxacin in the treatment of uncomplicated pyelonephritis. Am J Med. 1992;92(6A):86S–94S.

35. Opstrup KV, Bennike TB, Christiansen G, et al. Complement killing of clinical Klebsiella pneumoniae isolates is serum concentration dependent. Microbes Infect. 2023;25(4):105074.

36. West EE, Woodruff T, Fremeaux-Bacchi V, et al. Complement in human disease: approved and up-and-coming therapeutics. Lancet. 2024;403(10424):392–405.

37. Lukacsi S, Macsik-Valent B, Nagy-Balo Z, et al. Utilization of complement receptors in immune cell-microbe interaction. FEBS Lett. 2020;594(16):2695–2713.

